# Multi-view learning to unravel the different levels underlying hepatitis B vaccine response

**DOI:** 10.1101/2023.02.23.529670

**Authors:** Fabio Affaticati, Esther Bartholomeus, Kerry Mullan, Pierre Van Damme, Philippe Beutels, Benson Ogunjimi, Kris Laukens, Pieter Meysman

## Abstract

The immune system acts as the intricate apparatus dedicated to mounting a defence that ensures host survival from microbial threats. To engage this faceted immune response and provide protection against infectious diseases, vaccinations are the critical tool developed. However, vaccine responses are governed by levels that when interrogated separately only explain a fraction of the immune reaction. To address this knowledge gap, we conducted a feasibility study to determine if multi-view modelling can aid in gaining actionable insights on response markers shared across populations, capture the immune system diversity, and disentangle confounders. We thus sought to assess this multi-view modelling capacity on the responsiveness to Hepatitis B virus (HBV) vaccination.

Seroconversion to vaccine induced antibodies against HBV surface antigen (anti-HBs) in early-converters (n=21; <2 month) and late-converters (n=9; <6 months), was defined based on the anti-HBs titres (>10IU/L). The multi-view data encompassed bulk RNA-seq, CD4+ T cell parameters (including T-cell receptor data), flow cytometry data, and clinical metadata (including age and gender). The modelling included testing single-view and multi-view joint dimensionality reductions. Multi-view joint dimensionality reduction out-performed single-view methods in terms of area under curve and balanced accuracy, confirming an increase in predictive power to be gained. The interpretation of the findings showed that age, gender, inflammation-related gene sets and pre-existing vaccine specific T-cells were associated with vaccination responsiveness.

This multi-view dimensionality reduction approach complements the clinical seroconversion and all single modalities. Importantly, this modelling could identify what features predict HBV vaccine response. This methodology could be extended to other vaccination trials to identify key features regulating responsiveness.

## Background

The immune system is a complex network of interconnected components including effector cells, receptors, and mediators that underlie both humoral and cell-mediated immunity, innate and adaptive responses [1]. A single-level analysis of any of these components fails to fully capture the heterogeneity of the immune system and may not uncover meaningful markers. To address common research questions, a “systems immunology” approach is often necessary to leverage the interrelationships between different component levels. This framework will aid in detecting shared patterns and characterise the drivers of immune responses in a comprehensive manner [2]–[4]. Unfortunately, few methods are available within systems immunology that can directly interrogate immunological data across different levels.

Multi-view learning is a branch of machine learning techniques consisting in the fusion and integration of multiple datasets. By exploiting the additional intra-dataset information available, models thus built aim for a better generalisation performance [5]. Multi-view learning has received a lot of attention and several algorithms with encouraging performance have since been developed, with many based on Canonical Correlation Analysis (CCA) [6]. Multi-view learning has been successfully applied for biological data integration. In particular, CCA based applications have been implemented for oncological, Alzheimer’s studies and brain imaging tasks [7]–[11].

Principal Component Analysis (PCA) [12], [13] is a more well-known dimensionality reduction technique that computes a linear transformation of the original features to produce a new set of orthogonal features of lower dimensionality that capture maximal variability in the data. The derived variables are uncorrelated and are placed in descending order of their explained variance. CCA belongs to the same category of statistical techniques of PCA and it can be qualified as Joint Dimensionality Reduction (JDR). The main difference between the two methods lies in the input and the optimised statistic. While PCA operates on a single multivariate dataset at a time, CCA works on two datasets. CCA has since been modified to function on more than two datasets at once and exploit their interdependencies by maximising the sum of pairwise correlations (SUMCOR) [14]. The dimensions of the different views are simultaneously reduced while maximising correlation across modalities instead of variance, irrespectively of the data types.

The modelling for immunological datasets is unfortunately restricted to the common multi-omics data integration approach, which does not make use of CCA. This multi-omics method consists of a fine-tuned design aimed towards a specific set of data types, with Multi-Omics Factor Analysis (MOFA) as a prime example [15]. MOFA allows direct use of relational information to integrate the levels, but its specialisation locks their applicability to datasets of their field and hinders their use for immunology where other levels are more commonly measured, such as immune cellular composition or T cell and B cell receptor (TCR/BCR) repertoires.

Multi-view analysis is potentially beneficial to the vaccinology field. As highlighted in the literature, many factors influence the efficacy of a vaccination. These factors comprise identifying immune cells that are critical for a robust immune response including which memory cell population needs to develop (*e.g.* B cell vs T cells) and understanding how patient factors (*e.g.* gender, age, ethnicity) influence these findings. For instance, age significantly modifies the landscape of the immune system. These ageing changes are well documented in age-related pathologies such as Alzheimer’s, arthritis, hypertension and atherosclerosis, which are linked to chronic inflammation and interferon activation [16]–[18]. Additionally, pre-existing cross-reactive T-cells, such as in the CD4+ memory compartment, are required for a robust vaccine response to Hepatitis B virus (HBV)[19]. Therefore, multi-level factors need to be considered when determining who would respond best to a vaccination. However, the current applied approaches only analysed one layer at a time, and hence might have led to potential confounders in interpreting the data.

Among the unsolved questions in vaccinology is what drives and predicts a good immune response to a vaccine. Prior studies have shown that a vaccine response can in part be predicted using mRNA expression baseline levels from blood (cells) [20]–[22]. Thus, the response to a vaccine seems to be partially determined by the state of the immune system before the vaccine is administered. Predictive models based on a single data type, such as peripheral blood mononuclear cells (PBMCs), bulk transcription RNA expression, or flow cytometry (protein expression), have been previously built. However, as indicated before, these different measurement levels are not independent, with ∼40% of mRNA levels being correlated to protein expression [23]. Therefore, it is not known what is causative, and what is a related confounder. A multi-level approach may aid in deconvoluting these potentially confounding factors.

In this paper, we evaluate different multi-view approaches on multi-view data including bulk blood RNA sequencing (RNA-seq) gene expression, PBMC flow cytometry cell counts, memory CD4+ T-cell receptor repertoire and demographic metadata with reference to HBV vaccine response. We aimed to study whether this methodology can be helpful in determining the predictability of the baseline state for the vaccine response. As a proof of concept of our methodology, here we highlighted multi-view learning identified biological markers that correlate with response profiles developed after *de novo* HBV vaccination.

## Methods

### Study cohort

The study participants had been previously recruited to assess the HBV vaccination response [19], [24]. The cohort consisted of 34 HBV-naïve individuals, who were neither vaccinated nor infected (according to serology) against/by HBV. The original publications aimed to identify the response to a three-dose regimen of Engerix-B vaccine, with second and booster dose respectively administered 30 days and a year after the first dose. The vaccine response was measured based on anti-HBs titres captured by ELISA assays at days 0, 60, 180 and 365 (summarised in **Table 1**).

**Table 1.** Cut-off of converters and non-converters to hepatitis-B vaccination

| Outcome | Ab titre | Time | # in group |
| --- | --- | --- | --- |
| non-converters | <10IU/L | - | 4* |
| early-converters | >10IU/L | 60 days | 21 |
| late-converters | >10IU/L | 180 days | 9 |

\* Excluded from further analysis.

For the non-converter class, it is impossible to determine if seroconversion was eventually reached. For one non-converter, lack of space on the kit during TCR sequencing also precluded access to the related data modality. Non-converters were thus excluded from further analysis. Metadata of each participant were also considered (**Additional file 1**). All clinical features had been collected before administering the first dose. The features included age, gender, body temperature, maximum and minimum blood pressure (Min BP and Max_BP). The age distribution differed across all 3 classes (**Figure 1**), with late-converters tending to be older than the other groups (**Additional file 1**).

**Figure 1.**
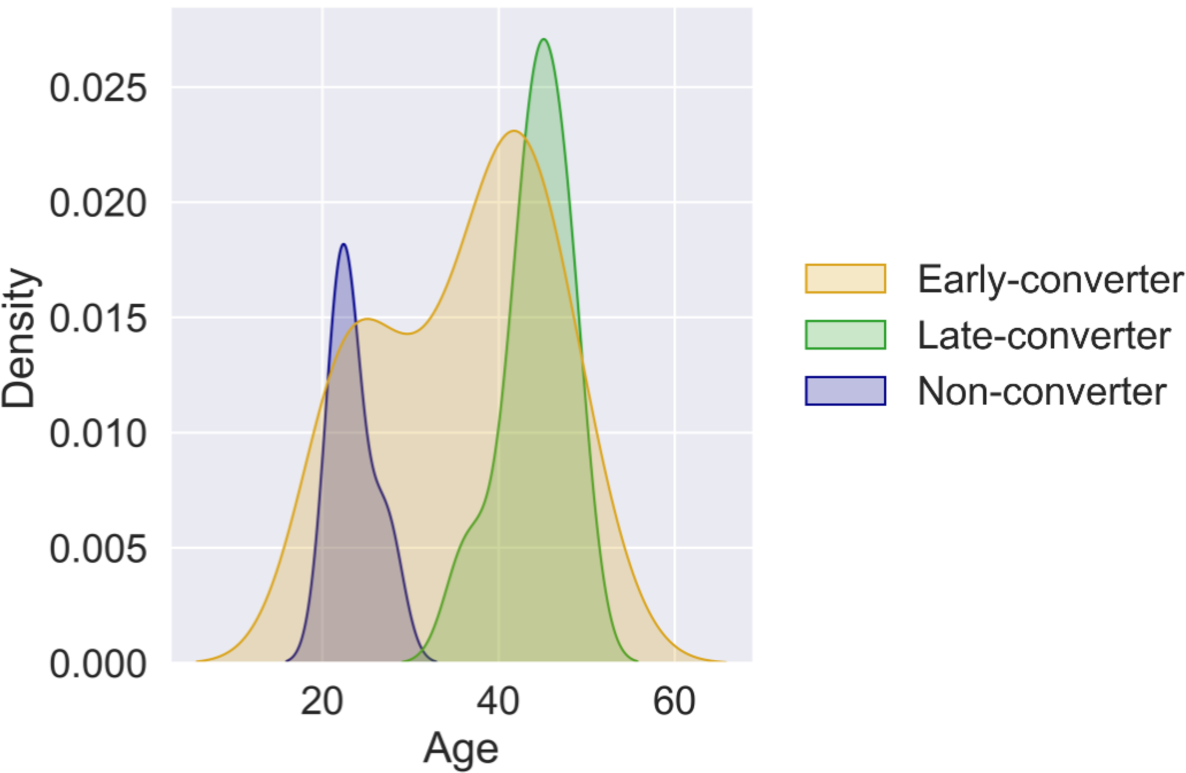
Age distributions reveal differences between classes. Age distributions per seroconversion class.

Other clinical factors available included access to the absolute numbers of white blood cells (with differentiation in monocytes, lymphocytes and granulocytes), red blood cells (with separate attributes for hemoglobins and hematocrit) and platelets were determined with a haematology analyzer [19], [24] (**Additional file 1**). Lastly, flow cytometry identified the percentages of CD4+ T cell relevant information (**Additional file 1**), as this population is critical to developing a robust HBV antibody response. These CD4+ T cell parameters included the normalised ratio of vaccine-specific T-cells (HepBTCR), the total number of TCRs sequenced in the CD4+ memory (B0), the frequency of bystander TCRs (PPnrB0) and the frequency of vaccine-specific TCRs (PSB0).

### RNA-seq data quality control steps

The first study [19] detailed the blood extraction and bulk RNA-sequencing of the PBMCs at day 0 and after vaccination at days three and seven. As stated, the pre-vaccination time point was considered. Of the 23812 transcripts, gender-specific and haemoglobin genes as well as removed transcripts with fewer than 100 counts across the samples were excluded, which left 13012 expressed transcripts for analysis. The counts were normalised using DEseq2’s median of ratios to correct for sequencing depth and RNA composition [25], to allow reliable between-samples comparisons. Differential gene expression analysis was performed with the DESeq2 negative binomial models. To improve the interpretability, the gene counts were aggregated based on the published BloodGenModule3 features to reduce the high dimensionality effects that this data view would carry in the analysis [26]. BloodGenModule3 was chosen, as it contained a hierarchical, fixed repertoire of 382 functionally annotated gene sets (blood transcriptional modules [BTMs]) [27]. The aggregation BTMs by means was used.

### Data sampling

The class distribution is relatively unbalanced. Synthetic Minority Oversampling TEchnique (SMOTE) [28] was used to upsample the minority class (late-conversion) in the training data set at each iteration of the cross validations used to estimate performance. It has been shown that SMOTE can be beneficial to high dimensional datasets when applied after feature reduction, such as in this case. New data points were generated by computing average measures of the five nearest neighbours of the same class in the training data set.

### Integration methods

The 30 early and late-converter samples, were used to evaluate several integration approaches for the applicability to the four processed views (**Additional file 1**). The workflow is detailed in **Figure 2**. Each data view was also initially inputted in the Logistic Regression (LR), multivariable model commonly used for biological sciences studies, to establish a baseline and examine the separate contributions. LR coefficients are readily interpreted, as the regression model requires few assumptions to determine the relationships between dependent and independent variables [29].

**Figure 2.**
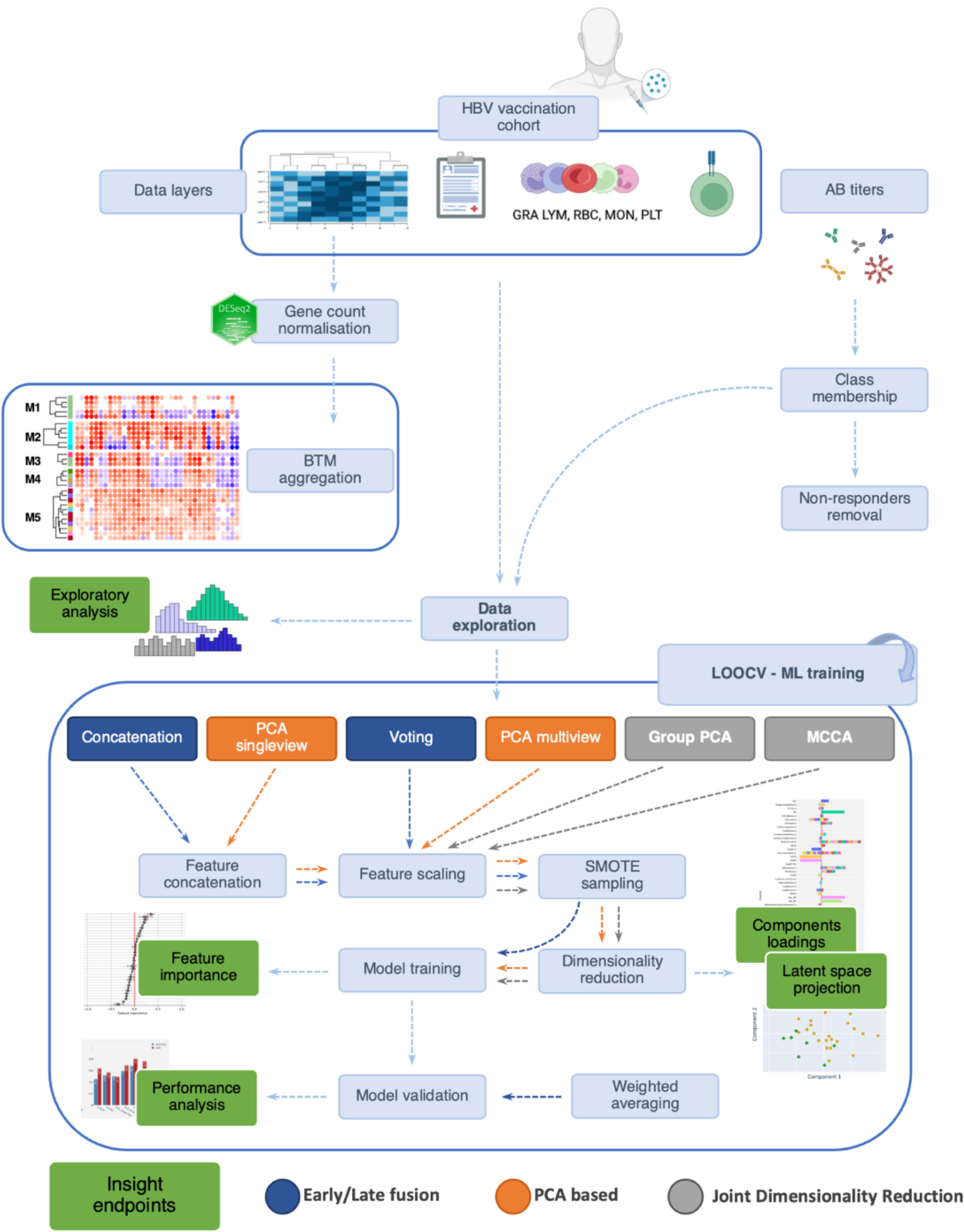
Research methodology workflow. The colour coding represents the integration philosophies (blue, orange and grey) and, in green, highlights the interpretability endpoints. Antibody (AB); Blood Transcription Module (BTM); Hepatitis B Virus (HBV); Leave-One-Out Cross-Validation (LOOCV); Machine Learning (ML); Multi-view Canonical Correlation Analysis (MCCA); Principle Component Analysis (PCA); Synthetic Minority Oversampling Technique (SMOTE). *Figure created using BioRender (BioRender.com)*.

The simplest and most common method to implement was an early fusion in which the views were concatenated at a feature level and a model is trained on all the variables. However, the advantageous ease of application is counterbalanced by the likely skewed fusion caused by high dimensional sets, such as our gene expression matrix [8], [30]. Mirroring this first technique a late fusion through voting was also applied [31]. The concatenation, in this case, happens at the decision level by merging independent predictions through weighted averaging. A classification model is built to exploit specific characteristics of the target view. The predictions are then weighted and combined in the final results.

Feature engineering was explored next. Variants of PCA were deemed most appropriate for our case. A sparse PCA variant [32], [33] that introduces a Least Absolute Shrinkage and Selection Operator penalisation [34] was applied on the concatenated features and its output fed as training to the models (named “PCA_singleview”). A PCA was also applied on each separate view to discover equivalent dimensions of the view-specific components. In case of the gene expression set the sparse variant was used. The embeddings obtained were then concatenated along the component dimension before training (named “PCA_multiview”). As an even more compact alternative each view was whitened with PCA, then, latent features were concatenated and a PCA was performed on this matrix once more (Group PCA) [35]).

An adapted multi-view version of CCA, MultiviewCCA (MCCA)[14], [35], [36] in its regularised form was implemented. The regularisation parameter was introduced to reduce overfitting caused by differences in dimensionality between datasets. Regularised MCCA aims to discover a common signal across sources while also reducing view-specific noise [37]. Sparsity was introduced to denoise gene expression in particular [38]. The number of features in this case greatly exceeds the number of observations and CCA could not be applied otherwise. Shared components can then potentially show meaningful discriminant features. Similarly to PCA, each component can be individually studied to reveal the factors that most contributed to it. This aspect is fundamental to draw valuable insights in the task at hand. The decision to reduce the summed dimensionality (399 features) of the multi-view data to a few dimensions allows to obtain substantially more interpretable results.

Some of CCA’s assumptions were considered during our analysis. Even though it was not strictly necessary for the machine learning algorithms used here, we applied standardisation on every dataset to respect CCA’s assumption of multivariate normal distribution and homogeneity of variance. The relationship between the canonical variates calculated and the original features is assumed to be linear unless a kernel variant is used. Similarly, to multivariate regression, and many other statistical algorithms, CCA gains inference power and reliability from a high sample size. The application of SMOTE thus contributed by generating more data points. These assumptions can also be applicable for every other PCA-based transformation we implemented.

## Results

### Anti-HBs titers separation

The anti-HBs conversion status was assessed based on the three time points after vaccination (**Figure 3A**). A PCA transformation of the antibody titer data revealed that principal component (PC) 1 (93% of the variation) segregates the data based on when the individuals responded to the vaccination. As expected, a clear separation occurred between the three clinically relevant groups: early converters, late converters and nonconverters (**Figure 3B**).

**Figure 3.**
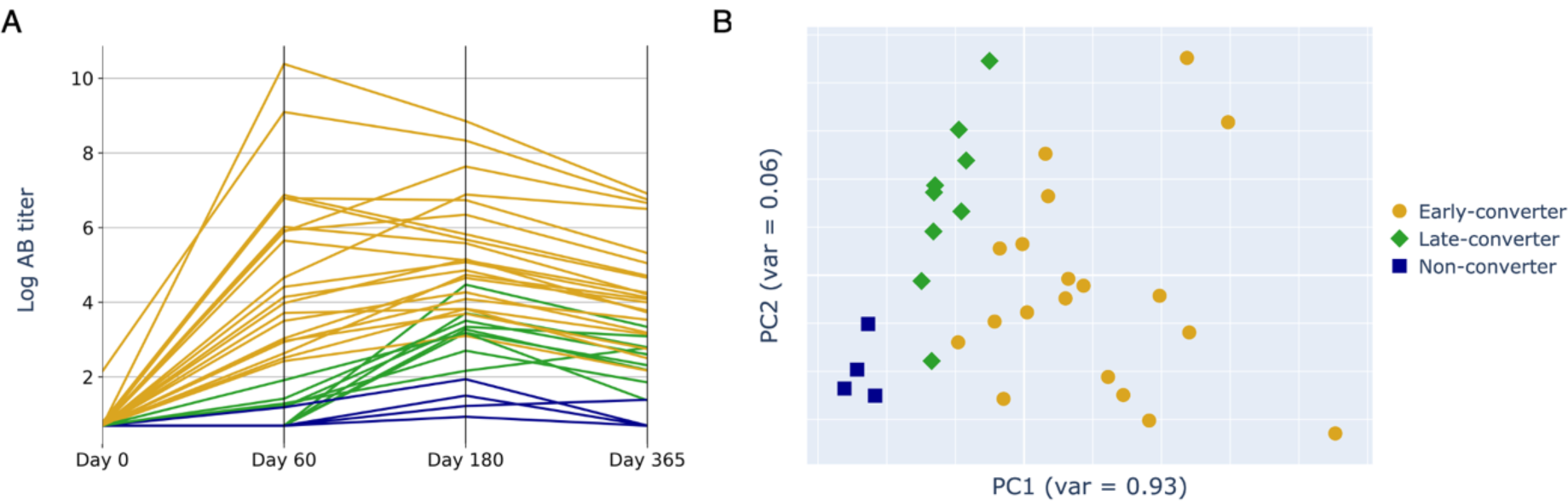
Principal components analysis of anti-HBs titres is correlated with responsiveness. (**A**) Antibody titers at different sampling time points on a logarithmic scale, divided across the three response classes. (**B**) Principal component (PC) 1 and PC2 of the antibody titer data. Coloured by converter type (early-converter = orange circle; later-converter = green diamond; non-converter = blue square).

### Independent data levels show correlated features

Collinearity across views is an expected characteristic within the multi-view HBV vaccine data set, as each molecular or clinical level will influence the others. This step was to uncover dependencies within the metadata irrespective of the vaccine response. As expected, several variables are strongly correlated (**Figure 4**). For example, the day 0 lymphocytes (LYM0) and granulocytes (GRA0) counts are strongly anti-correlated (R = −0.98). As per the literature [38], the expected negative correlation of female gender (Gender_F) and red blood cell count (RBC0) (R=-0.63) was identified. TCR-derived parameters were also assessed. The day 0 total unique TCR sequences per individual (B0) is correlated with both the fraction of HBs-Ag-specific (PSB0, R=0.88) and aspecific (PPnrB0, R = 0.61). The normalised ratio of vaccine-specific TCRs (HepBTCRs) was poorly correlated to the initial TCR counts (B0; R = 0.16).

**Figure 4.**
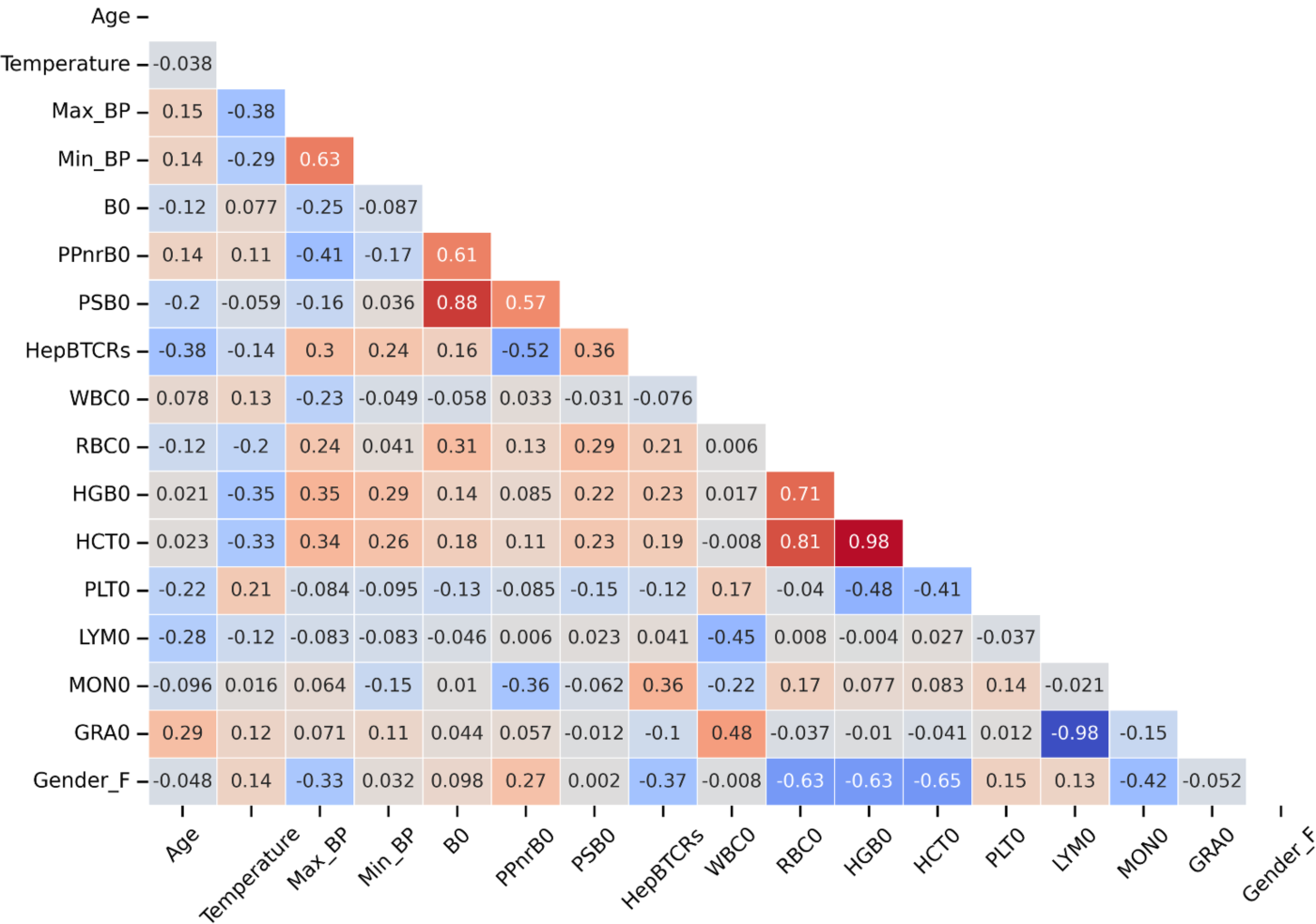
Correlation matrix shows overlap of cohort features. Data represents the correlation coefficient across each non-genetic feature (see **Additional file 1** for cohort details).

### Individual levels contain elements that correlate with response

Before implementing the multi-view integration, we interrogated the data to identify key features. Foreseeably, *a*ge is higher for late-converters (p-value = 0.019, **Additional file 2**). Age is also anti-correlated to the anti-HBs titers presented at each timepoint, especially after the first two doses (**Additional file 3**). Within the flow cytometry modality, lymphocyte counts seem to possess a positive association with the early response (**Additional file 1**). On the other hand, white blood cells and in particular granulocytes are associated with a delayed response (**Additional file 4**). However, possibly due to outliers, no strong statistical significance emerges from these comparisons. Conversely, testing of the ratio HepBTCR reveals a significant association with an early HBV vaccine response traceable to the higher HBV specificity of TCRs (p-value = 0.003, **Additional file 5**). Just one gene was identified as differentially expressed prior to vaccination when comparing the early and late-converters: HLA-DRB5 (Log_2_FoldChange = −4.96, adj. P-value = 2.50e-4 [False discovery rate = 0.05]). HLA-DRB5 was down-regulated in the late-converters, and encodes a part of the MHC class II complex, related to antigen presentation and CD4+ T cells.

### Projection of data views provides insights into response classes

The different data levels were integrated and projected using two distinct approaches (**Figure 5**). The first is a PCA, where the PCs are defined by those that capture the most variance across the entire, concatenated, dataset. The first two PCs can be evaluated through visualisation of the samples in the lower dimensionality latent space (**Figure 5A**). The second is MCCA, which aims to detect a common latent structure across multiple datasets by following the SUMCOR definition. The two most informative canonical variates after same-time MCCA integration can then be inspected (**Figure 5B**). The MCCA approach had more prominent separation in the data.

**Figure 5.**
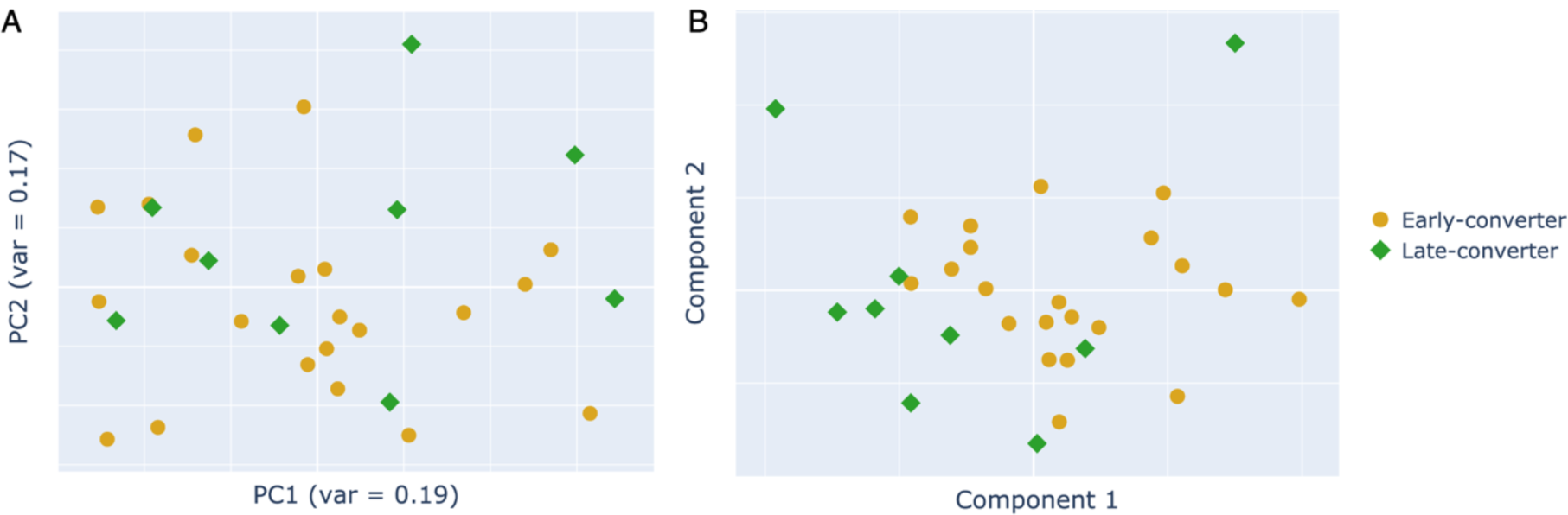
MCCA latent space projection provides superior class separation. Two-dimensional data projection with (**A**) PCA and (**B**) MCCA of the integrated multi-view vaccine dataset. Each scatter plot showcases the first two dimensions. Data represents the early-converters (orange circle) and late-converters (green diamond) after applying each model. Multi-view Canonical Correlation Analysis (MCCA); Principal Component (PC); Principal Component Analysis (PCA).

As MCCA has better separation of the converter status, it can further be explored by considering the loadings underlying the new dimensions which represent the contribution of each base feature. **Additional file 6** shows the stacked coefficients for each of the two first canonical variates of MCCA, grouped by function. These effects were attenuated if GroupPCA was considered (**Additional file 7)**.

There are certain pre-existing features that are associated with vaccine response [19], which include specific gene expression patterns (*e.g.* cytokine expression), immune cells overrepresentation and patient factors (*e.g.* age, gender, comorbidities). Upon inspection of the first component of the MCCA projection age, oxidative stress, inflammation and interferon modules all have negative contribution (**Additional file 6A**). Lymphocytic BTMs present a negative sign to their coefficients as well. Furthermore, erythroid cells modules and blood cells related counts are relevant for both dimensions and in particular for the second component (**Additional file 6B**). As can be expected based on the prior pairwise correlations, higher red blood cell counts are also associated with the male gender parameter [39]. The signal given by monocytes is weaker, but more consistent for the second component, with the monocyte count and monocyte-related modules having a positive contribution. Finally, CD4 lymphocytes are present in the contribution for the two components both in terms of modules and counts coming from the flow cytometry assays. This compound observation reinforces the importance of lymphocytes for the projection.

Moving to GroupPCA, we can identify, for the first PC (**Additional file 7A**), several age-related markers. In the present study, inflammation, interferon, oxidative stress and phosphorylation signatures appeared positively tied to ageing. Other positive interactions included higher min and max blood pressure and a higher cell death-driven gene expression as well as lower red blood cells counts (RBC), hematocrit (HCT) and haemoglobin protein counts (HGB). Finally, the erythroid cells BTMs expression is noticeably relevant for this component. Platelets counts and BTMs appear negatively related to age.

For the second PC (**Additional file 7B**), at lower ages we also see lower blood pressure, reduced cell death, oxidative stress and phosphorylation. Diminished T cells and lymphocyte related features seem to signal a less active adaptive immune system. On the other hand, heightened white blood cells counts, neutrophils and neutrophils activation modules, granulocytes counts together with the presence of inflammation modules still show innate immunity characteristics for this dimension.

In comparison to MCCA, GroupPCA revealed distinct modules (**Additional file 7**). The top homogenous contributions included interferon and inflammation modules. This was distinct from MCCA (**Additional file 6**) where these features were not represented as having coherent contributions. GroupPCA presented a stronger T cell signal. Both methods assigned rather consistent importance to gene transcription, protein synthesis and protein modification modules for both of their dimensions.

Additionally, to complement the previous analysis and to disentangle loadings that present conflicting contributions, features significantly correlated to one of the latent components (p-value < 0.05) can be overlaid on top of the MCCA projected cohort. This visualisation allows to identify a cohesive inflammatory signature. A region of the latent space is thus univocally characterised by elevated temperature, age, granulocyte counts, inflammation and neutrophil related gene sets (**Figure 6B**). As a result of the inferior class separation, only a more ambiguous sub-setting of the cohort can be derived from visualising correlations of the same features on the standard PCA (**Figure 6A**).

**Figure 6.**
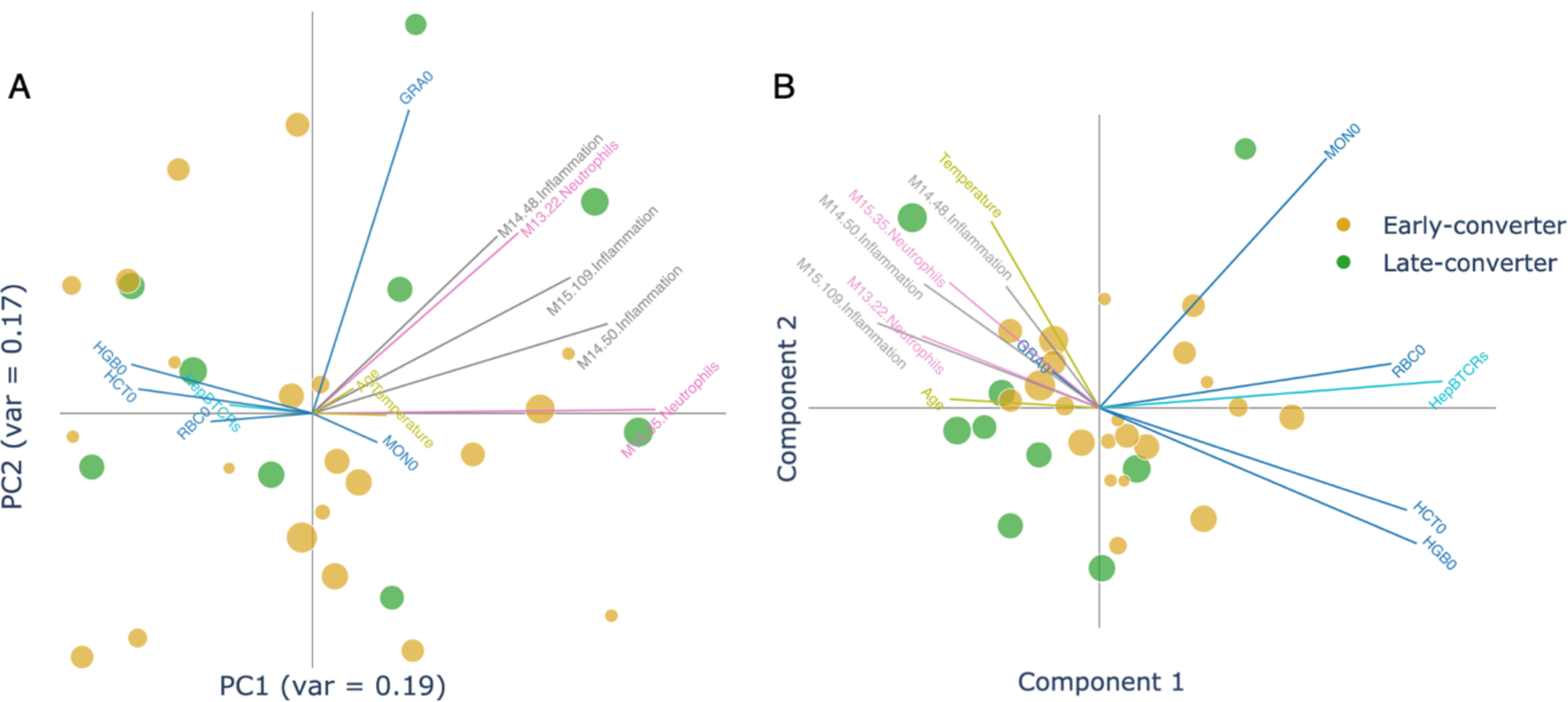
MCCA latent space projection shows inflammatory markers characterising part of the cohort. Two-dimensional data projection with (**A**) PCA and (**B**) MCCA after the application of a feature correlation overlay. Features significantly correlated to the MCCA projected space reveal a distinct inflammation-associated region. Marker sizes are proportional to the age of the individual. Feature Vector colours identify separate views and module functions. Vector coordinates are calculated using the Person’s correlation to the respective latent component. Granulocytes count (day 0) (GRA0); hematocrit (day 0) (HCT0); normalised ratio of vaccine-specific TCRs (day 0) (HepBTCRs); haemoglobin protein count (day 0) (HGB0); Multi-view Canonical Correlation Analysis (MCCA); Monocytes count (day 0) (MON0); Principal Component (PC); Principal Component Analysis (PCA); red blood cells count (day 0) (RBC0).

### Multi-view integration allows superior classification performance

Thus far, the integration of this data set has been explored in an unsupervised fashion, *i.e.* without prior knowledge of the converter classes. However, as these are known, the various multi-view integration methods can be evaluated in their ability to capture and predict this ground truth. This can be done by feeding the integration output into a supervised machine learning model with a Leave-One-Out-Cross-Validation (LOOCV) approach. Integration approaches that successfully capture latent patterns within the data set that are related to vaccine response, should be able to obtain a high performance.

The LOOCV was repeated 20 times with random initialisations to identify a more reliable performance spread (**Figure 7**). The classification task was originally skewed with a majority class of 21 early-converters versus a minority class of 9 late-converters. Evaluation of each data layer independently with a LR (**Additional file 8**), revealed that not all layers held the same predictive capacity. Overall, the TCR layer had the strongest predictive performance on validation data, with both balanced accuracy and area under the receiver operating characteristic curve (AUC) measures above 0.7 (**0.708±0.067** accuracy; **0.731±0.040** AUC). Conversely, the outputs produced using cell counts or metadata were slightly better than a random classifier, corresponding to a prediction accuracy or AUC marginally above the threshold of 0.5. The model using gene expression data consistently assigns a prediction with confidence (>0.6 decision value; **Additional file 9**). This results in a higher AUC (**0.611±0.009**) but a lower balanced accuracy (**0.440±0.018**).

**Figure 7.**
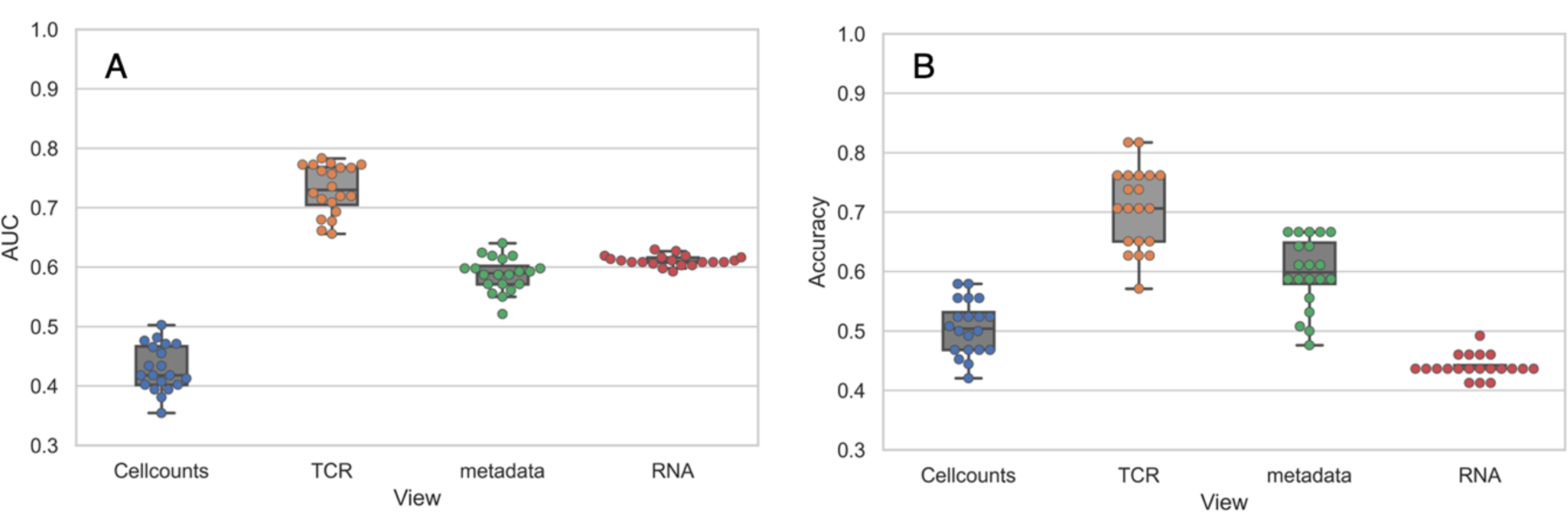
Single view performance comparison shows the superior predictive power of the TCR-seq layer. Methods performance comparison for the Logistic Regression classifiers trained on single modality data in terms of (**A**) AUC and (**B**) accuracy. Area under the receiver operating characteristic curve (AUC); bulk blood ribonucleic acid sequencing data (RNA); T-Cell Receptor sequencing data (TCR); T-Cell Receptor Sequencing (TCR-seq).

To boost the performance of the models, joint dimensionality reduction algorithms were applied. As expected, it improved the LR fitting for both GroupPCA (**0.750±0.034** AUC; **0.655±0.051** accuracy) and MCCA (**0.731±0.042** AUC; **0.627±0.047** accuracy) (**Figure 8A**). GroupPCA surpassed both the single level baselines and the early fusion, while also adding the interpretability highlighted in the previous section. However, using regular PCA methods was not as effective or even counterproductive, causing, in some instances, a drop in performance compared to the best single-view predictors (**Figure 8A**).

**Figure 8.**
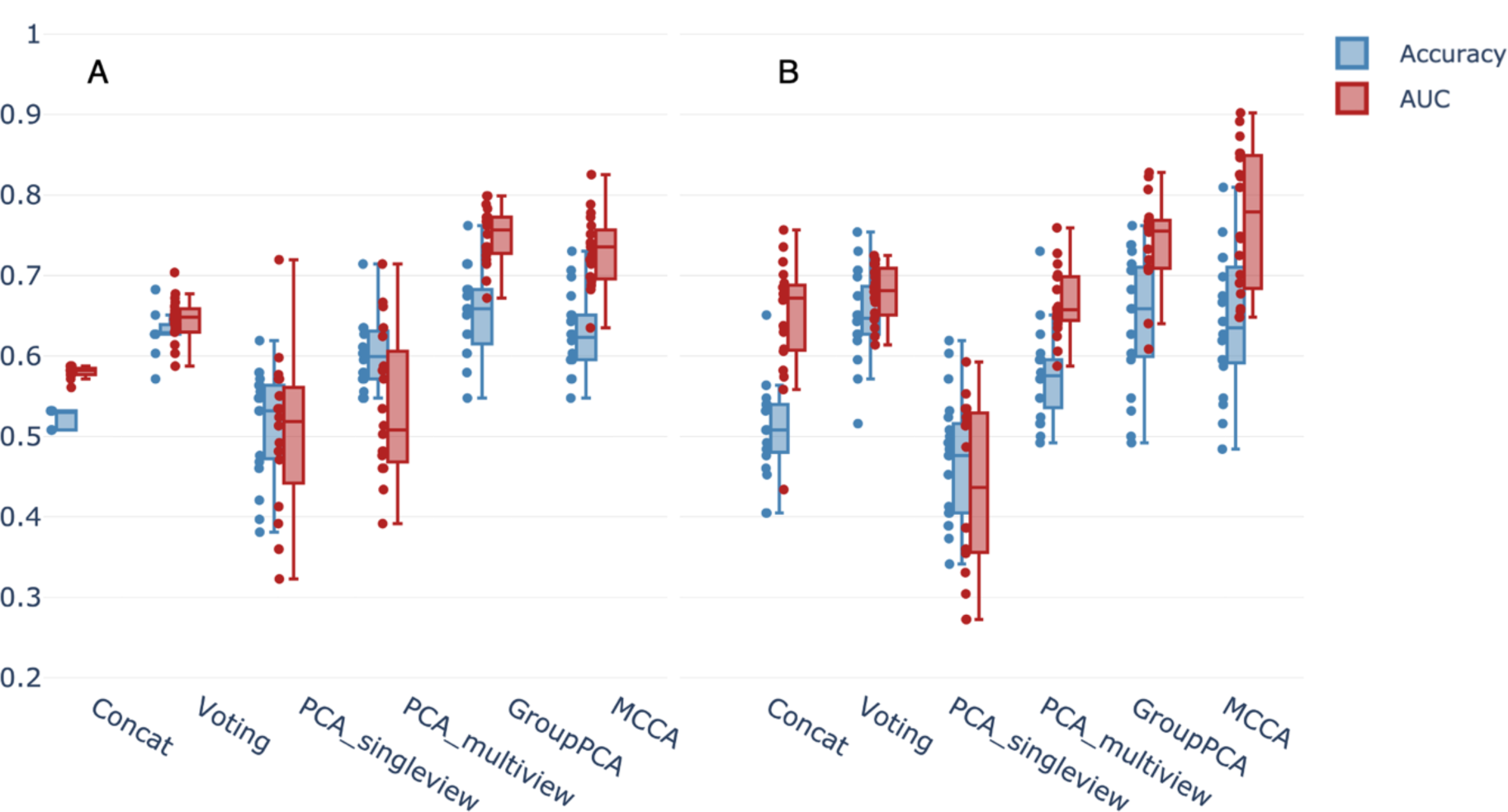
Classifiers performance comparison shows positive impact of joint data integration on predictions. Methods performance comparison for the (**A**) Logistic Regression and (**B**) Random Forest classifiers in terms of AUC and Accuracy. Area under the receiver operating characteristic curve (AUC); *Multi-view Canonical Correlation Analysis (MCCA); Principal Component Analysis (PCA)*.

The data integration techniques produced consistent results for the more complex model, Random Forest (RF) (**Figure 8B**). Once again and alongside GroupPCA, MCCA presents itself as the best integration method. JDR implementations fell in the range of 0.64-0.8 for both AUC (**0.743±0.053** GroupPCA; **0.770±0.087** MCCA) and accuracy (**0.646±0.079** GroupPCA; **0.640±0.082** MCCA), performing comparatively well, albeit presenting a larger spread. In this setting PCA single view was the only method to present a poor performance. After such a strong dimensionality reduction this result can be expected. Models are effectively trained on very few features and a simpler method should be sufficient in this circumstance.

In addition, RF and LR can be easily unravelled to further allow for interpretability of the underlying predictions. The coefficients of the LOOCV LR trained on single data views (**Figure 9**). Old age was the leading factor for a late response (negative class) in the unimodal analysis (**Figure 9C**), which was also apparent in our previous modelling.

**Figure 9.**
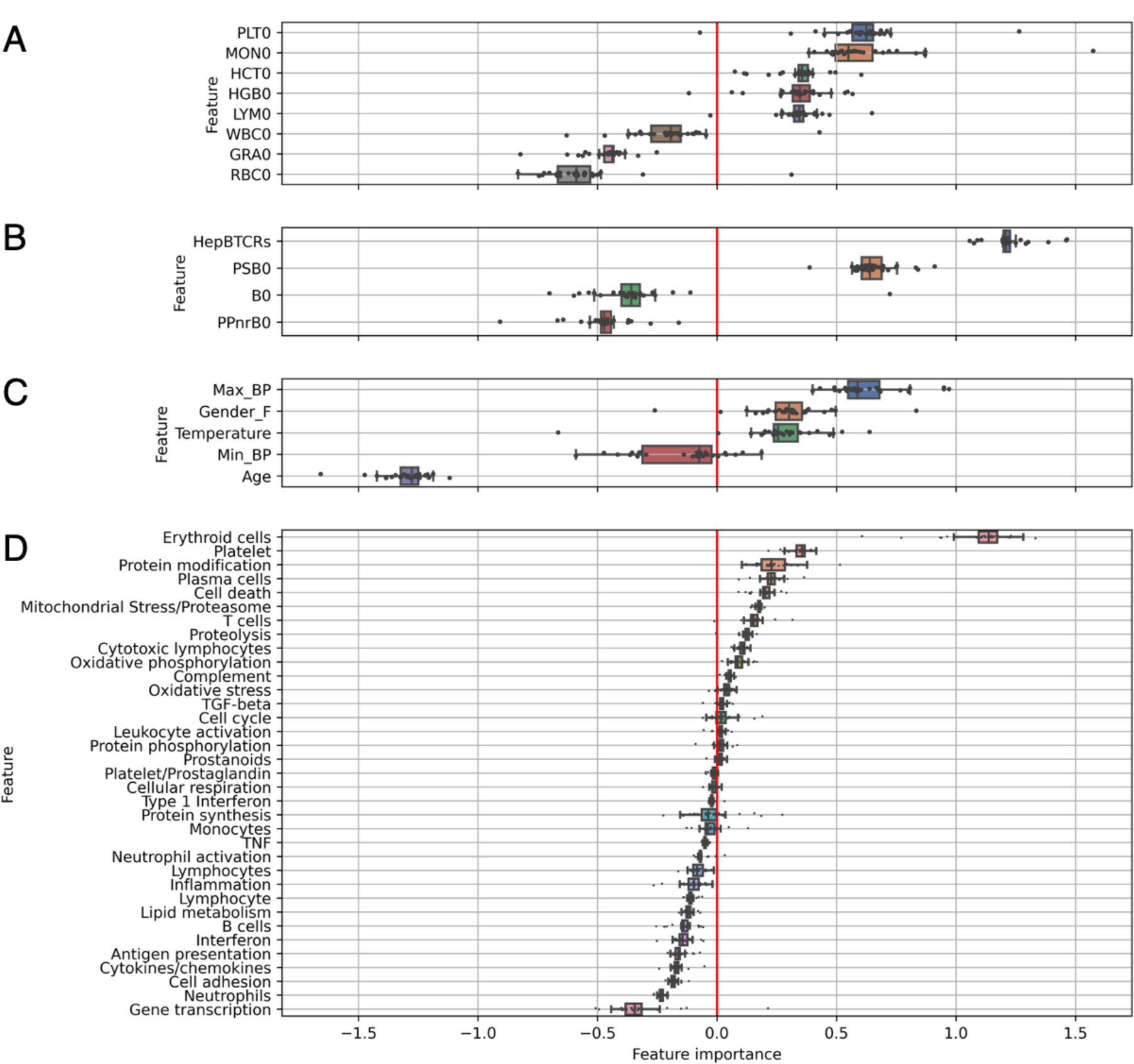
Unimodal feature importance analysis bolsters insights into biological markers tied to vaccine responsiveness. Importance of leave-one-out cross validated LR coefficients for the unimodal models: (**A**) Cell counts, (**B**) CD4+ T-cells, (**C**) Metadata, (**D**) BTMs. Positive coefficients favour early-response predictions while negative coefficients favour late-response predictions. The vertical red line indicates the zero-importance threshold. BTMs with unknown functionality have been excluded from the visualisation. Blood Transcription Module (BTM); Logistic Regression (LR).

Here feature importance within the dataset is showcased, which included granulocyte counts, neutrophils modules and other immune related BTMs linked with late response predictions. These features were distinct from the earlier converters which included the female gender, temperature, pre-existing vaccine specific T-cells (**Figure 9B**), platelets counts and BTMs, monocytes counts as well as erythroid cells associated counts and modules. The coefficients for platelet counts, erythroid cells and platelet BTMs, present the highest median for the single-view flow cytometry and RNA-seq models respectively (**Figure 9A and Figure 9D**).

## Discussion

Here we showcased the utility of using multi-view analysis to aid in the interoperability of a longitudinal dataset following response to HBV vaccination. This analysis involved incorporating anti-HBs titre levels, RNA-sequencing, CD4+ T cell-receptor, flow cytometry and patient defined features (*e.g*. age, gender). This multi-view analysis implementation aided in finding interpretable patterns in a vaccination context, which has not been previously interrogated together. This study bases itself on the notion that prior studies have also highlighted relations between different levels (*e.g.* mRNA expression, CD4+ T-cells, age and BMI) and the eventual HBV vaccine response, as measured by the anti-HBs titers [19], [24], [40], [41]. To point out the most meaningful approach and improvement of predictive power of multi-view datasets, we used several integration methods. Our approach identified key baseline features associated with earlier and later converters after HBV vaccination. This modelling was able to make informed interpretations of the data and account for confounding factors. For instance, our modelling was able to highlight that older age and B-cells were more associated with late-converters.

Age is undoubtedly a major factor in the immune response, especially with “inflammaging” and “immunosenescence” being well documented phenomena [16], [17], [42]. The impact of age is so broad that, for instance, even though monocyte numbers have not been shown to differ in older populations when compared to younger subjects, their phenotype and functionality is considerably different with higher degree of senescent qualities [43]. Inflammation, interferon, oxidative stress and phosphorylation dysfunctions in elderly are also widely confirmed by the literature [44]. In our study, these signatures appear along with higher min and max blood pressure and a higher cell death-driven gene expression. Lower RBC, HCT and HGB in conjunction with the positive contribution of age, let us infer these patients had an ageing anaemia and reduced hemoglobins [45].

Granulocytes and lymphocytes appeared mutually exclusive, showing a very strong anti-correlation, a behaviour also detected in the literature [46]. According to [46]–[48], a high neutrophil(granulocytes)/lymphocyte ratio is tied to the inflammatory status of the patient and to an impaired cell-mediated immunity. Conversely, a lower ratio is associated with better outcomes for several diseases, including HBV. Likewise, higher temperatures can have a limiting effect on viral replication and play an enhancing role in many immune related functions [49]. Our model contextually showed both patterns as important for conversion classification.

Gender presented itself as another important factor for the interpretation of the results. The male gender, relevant in this study, has previously been shown to influence the presence of pro-inflammatory markers [50]. Additionally, females have been documented to have a better response to HBV booster vaccination with greater anti-HBs titers [51]. The models presented here showed the male gender was associated with inflammation-related features and predictive of early response when controlling for other variables during regression.

For the early response predictions, and in accordance with results for other pathogens [52], [53], pre-existing vaccine specific TCRs contributed positively. Our analysis, through T-cell gene expression and CD4 counts contributions, highlights potential existence of pre-vaccination priming of the T-cell compartment that may be conducive of an early vaccination response, and requires follow-up interrogations on which CD4+ T cells contribute to vaccine response. Using this modelling approach, we could also mine that erythroid cells and platelets counts as well as their respective BTMs were also impactful for an early seroconversion. Intriguingly, platelets counts and BTMs were negatively correlated with age and had opposite contributions on the single-view model predictions, in a behaviour confirmed by the literature [54]. This data can be interpreted as a reduced number of immune cells simultaneously circulating in the blood, leading to an overall less activated system.

Among the dimensionality reduction techniques, GroupPCA was on par with MCCA, surpassing every other method and all four of the single level models in terms of AUC. However, fine tuning of the regularisation or the addition of learned weighting parameters to MCCA’s algorithms can boost its performance. Furthermore, MCCA seems to have found a parsimonious representation of heterogeneity in the sample (significant latent space). The separation it produces was more prominent and, we can assume, more trustworthy than single-level labelling.

Our modelling achieved a reasonable level of accuracy and AUC, but did not reach a median AUC>0.8, which highlights that improvements could be made with further refinements. For instance, a parameter of “importance” can be learned through self-attention mechanisms for each view to weight the contributions differently when applying JDR, instead of applying an imposed regularisation parameter. Starting from the assumption that similarity between samples is different in different views, [55] proposed a smooth representation by using a self-weighting method that enhances similarity grouping effects. An explanation for the reduced AUC could be attributed to incorrect converter classification, as this was based solely on the AB titres. However, our data identified screening for memory CD4+ T cells would identify if someone responded to the vaccination. Additionally, due to the limited understanding of all possible gene-gene networks (gene modules), this study’s modelling could not map the contribution of genes with unknown biological function. Therefore, alternative ways to identify the most relevant gene sets is needed. Future versions of the method will have the capacity to include more gene modules, other than [27] and allow to use first dimension PCA instead of the BTM mean to allow capturing dataset-specific variability.

To solve the linearity constraint of the combinations, Kernel MCCA (KMCCA) [35] could be used to first project the data in a higher dimensional space via the kernel trick and then apply MCCA to the new feature space. Alternatively, similarity network fusion (SNF) [56] has emerged as a specific data integration technique to learn underlying global and local structures when dealing with patient data [57], [58]. Producing embeddings, with encoders, could help us handle missing values, missing views and not only that, help us even reconstruct these missing aspects. In this field where data is scarce and missing data can skew the findings. At training time all available views can be used to learn an embedding. During testing or generalisation, the embedding can help reconstruct the missing views’ information since it was learned and it was incorporated in the embedding. Successful application of Deep CCA and Deep Canonically Correlated AutoEncoders (DCCAE) [59] prove the feasibility. Although these implementations are promising, deep learning models had to be excluded at the time of this study, due to the limited sample size.

For the purpose of the analysis, a few drawbacks have to be considered. Such as the non-cohesive nature of the contributions coming from the BTMs. Firstly, the noise of gene expression data hinders full readability of their contributions. The high dimensionality aspect of RNA-seq data made the pattern harder to interpret. However, noisy results were expected when dealing with such noisy data. This problem is not unique to this study but is widely understood to be an unsolved challenge. Secondly, an effect akin to Simpson’s paradox [60], [61] caused by multicollinearity, presents itself when inspecting some of MCCA’s loadings. For example, HCT, RBC and HGB are strongly positively correlated and we would expect them to appear with same-sign coefficients in our analysis. Yet, the sign of the relationship is reversed. CD4 lymphocytes are present in the contribution for the two components both in terms of modules and counts coming from the flow cytometry assays, but the aforementioned effect limits their unambiguous interpretation. GroupPCA seemingly introduced less of this inversion paradox. On the other hand, it was possible to address this issue by visualising the correlations between the input features and the projected dimensions. The true relationships between variables could hence be revealed.

Here our multi-view analysis identified a dynamic and multi-faceted HBV immune response linked to many factors for early and late responders. When designing this type of experiment, an important aspect to consider is if the response is primary T cell vs B cell response, as some vaccines show primarily a T-cell based response [62], [63]. Furthermore, when designing this type of experiment, one needs to consider the most approximate time points, which may be not be applicable to the different kinetics (*e.g.* mRNA, live attenuated, Toxoid, viral vector, etc..). [64]. The types of primary responder cell will change which time points to collect. Currently, most attempts to predict vaccine immune responses are based on the sole antibody titers. This measurement tends to disregard the heterogeneity of the adaptive immune response and disregard the T cell contribution. With the unsupervised approach, JDR has been shown to help uncover a space in which separation between classes of interest is more pronounced. However, the findings are currently restricted to HBV vaccine, and testing for other vaccines is needed. We effectively dealt with an unbalanced phase 1 trial-sized population for this analysis. The feature engineering and data integration steps had a decisive effect both in terms of performance and are paramount to gain actionable insights. Ideally a bigger sample should be used for the validation setting.

### Conclusion

Our multi-view modelling reflected many aspects of what is known about natural HBV infections. Our findings on vaccination data, mirror HBV signatures present in the literature. The results presented throughout this paper suggest the methodology is valid to identify biologically important features and why multi-level interrogations can aid in finding the most critical features. This study’s modelling approach identified that early converters have more in common with those that respond well to the natural infection, while late responders were associated with older individuals and an inflammatory phenotype. Thereby, this methodology could recapitulate the known finding in the literature regarding HBV response. To confirm the utility of this modelling, we recommend testing the modelling on other HBV vaccination cohorts or in other immunological settings. We envisioned this type of framework will become useful for guiding rational vaccine antigen discovery as well as for the use of targeted adjuvants, along with, many other applications.

## Declarations

### Availability of data and materials

The datasets analysed during the current study and the code used to generate the results are available on GitHub at https://github.com/fabio-affaticati/MultiModHepB.

### Competing interests

KL and PM hold shares in ImmuneWatch BV, an immunoinformatics company.

## Funding

This work was supported by the Flemish Government under the “Onderzoeksprogramma Artificiële Intelligentie (AI) Vlaanderen” Program, the University of Antwerp (BOF GOA: PIMS), the European Research Council (grant agreement 851752-CELLULO-EPI) and Chan Zuckerberg Initiative (STEGO).

## Authors’ contributions

FA performed the study and wrote the manuscript. EB performed the original study on hepatitis B vaccine response. KM supervised and revised the manuscript. KL and PM conceived and supervised the study and revised the manuscript. PVD, PB and BO revised the manuscript. All authors read and approved the final manuscript.

## Acknowledgements

George Elias, Geert Mortier, Arvid Suls, Viggo Van Tenderloo & Hilde Jansens.

## Notes

### Summary of Updates

Abstract rephrasing; some figures have been reordered ; added Figure 6 and 7; Figure 8 improved visually; updated declarations; general polishing in the writing of some sections.

