## Additional files for "Multi-view learning to unravel the different levels underlying hepatitis B vaccine response"

***Additional file 1. Comparison table of the baseline non-genetic attributes between early and late converters.***

| Dataset | Feature | Early-converters (n=21) average (range) | Late-converters (n=9) average (range) |
|---|---|---|---|
| Metadata | Age (years) | 35.9 (21.3-50.2) | 44.19 (36.3-48.5) |
| | Temperature ([degree]C) | 36.60 (35.92-37.25) | 36.57 (35.35-37.19) |
| | Max BP (mmHg) | 126.90 (98-148) | 122.11 (103-157) |
| | Min BP (mmHg) | 75.62 (55-93) | 74.67 (59-105) |
| | Female/Male | 13/8* | 7/2* |
| CD4+ T cell parameters | B0 (counts) | 50790.38 (37866-71140) | 50237.89 (29330-87551) |
| | PPnrB0 (%) | 2.71 (2.01-3.45) | 3.37 (1.72-5.62) |
| | PSB0 (%) | 2.75 (1.87-3.87) | 2.49 (1.40%-4.53%) |
| | HepBTCRs (counts) | 1.02 (0.74-1.36) | 0.78 (0.42-1.11) |
| Cell counts | WBC0 (counts) | 6.10 (3.6-9.4) | 6.74 (4.3-9.3) |
| | RBC0 (counts) | 4.59 (3.94-5.21) | 4.60 (4.14-5.42) |
| | HGB0 (counts) | 13.62 (11.5-15.5) | 13.30 (8.1-15.4) |
| | HCT0 (counts) | 40.67 (35.0-46.4) | 39.82 (26.2-46.2) |
| | PLT0 (counts) | 243.67 (144.0-351.0) | 242.89 (172.0-403.0) |
| | LYM0 (counts) | 34.14 (19.5-47.8) | 29.7 (15.3-38.7) |
| | MON0 (counts) | 4.79 (3.2-8.2) | 4.13 (2.5-7.4) |
| | GRA0 (counts) | 61.07 (49.0-76.6) | 66.17 (57.4-77.3) |

*See the original manuscripts* [19], [24] *for breakdown of features.*
*Abbreviations: Max BP = systolic blood pressure (day 0), Min BP = diastolic blood pressure (day 0), B0 = total unique TCR sequences (day 0), PPnrB0 = fraction of HBs-Ag-aspecific (day 0), PSB0 = fraction of HBs-Ag-specific (day 0), HepBTCRs = normalised ratio of vaccine-specific TCRs (day 0), WBCO = white blood cells count (day 0), RBC0 = red blood cells count (day 0), HGB0 = haemoglobin protein count (day 0), HCT0 = hematocrit (day 0), PLT0 = platelets count (day 0), LYM0 = lymphocytes count (day 0), MON0 = monocytes count (day 0), GRA0 = granulocytes count (day 0).*
**Number of individuals.*

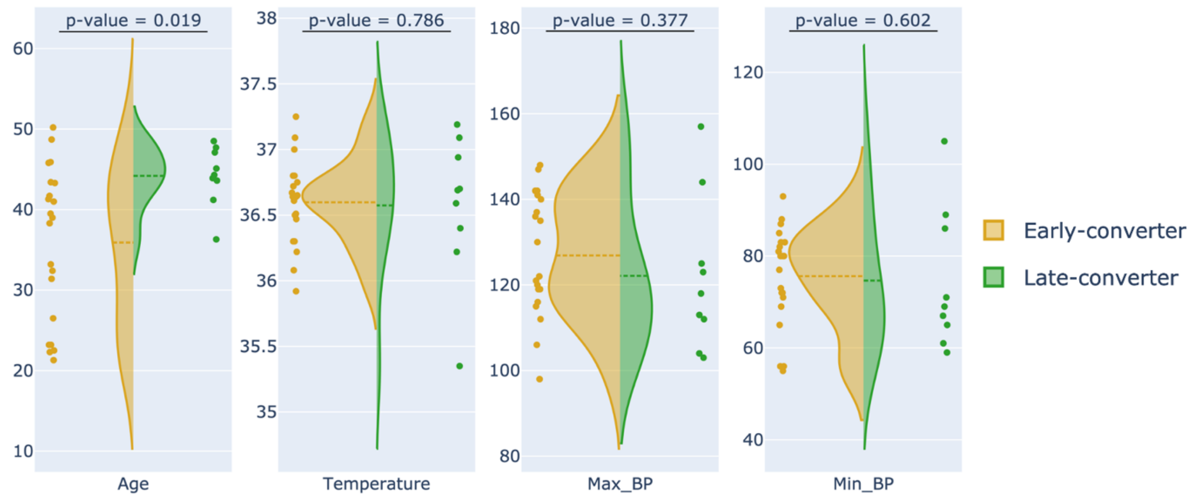


***Additional file 2. Metadata distributions reveal age differences.*** *Metadata feature distributions per class. Age is significantly higher for late converters (p-value = 0.019, two-sided Mann-Whitney U-test).*

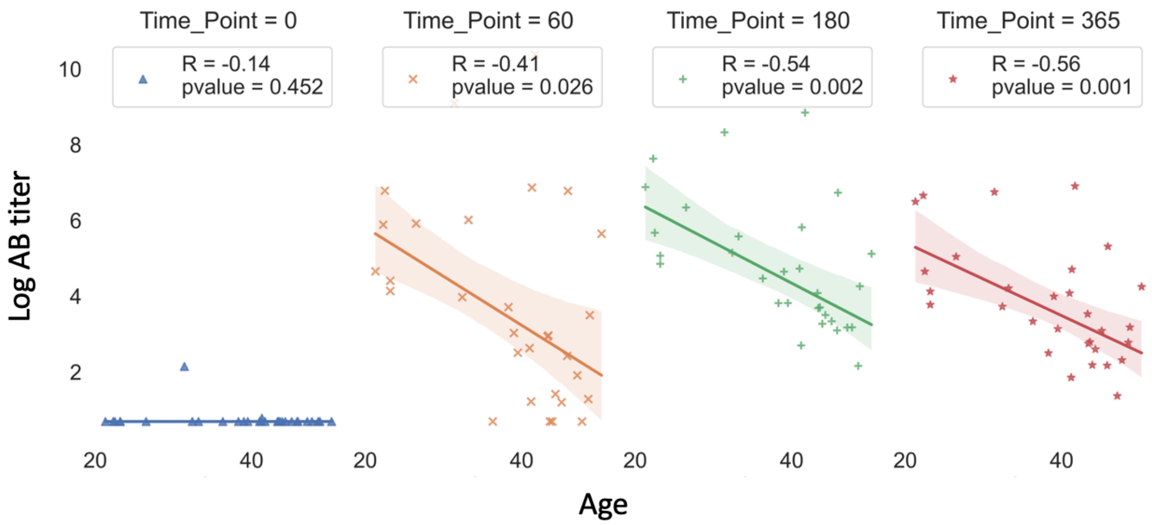


***Additional file 3. Longitudinal analysis of AB titers shows anticorrelation with age.*** *Comparison between age and AB for early and late converters, presented at each timepoint of the study. Time is expressed in days from the first vaccine shot.*
*Antibody (AB).*

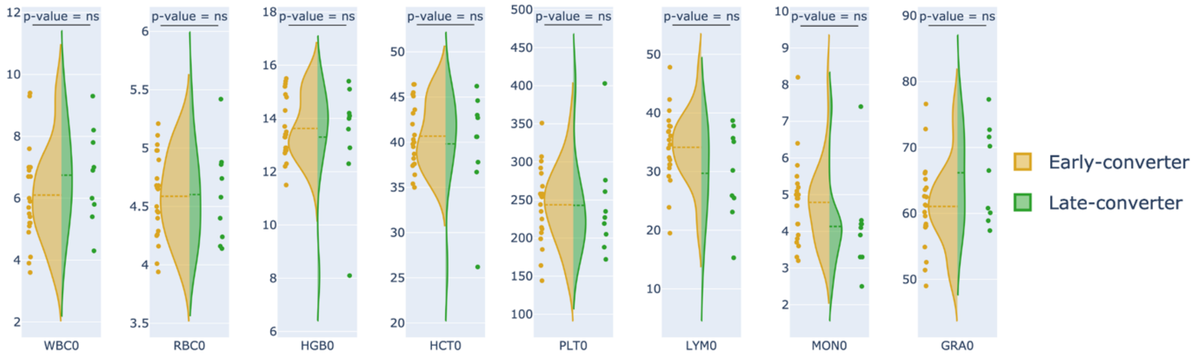


***Additional file 4. Cell counts distributions reveal class differences.*** *Cell counts feature distributions per class. Granulocytes and monocytes appear in lower quantities, respectively, for early and late converters. However, possible outliers lead to non-significant differences.*

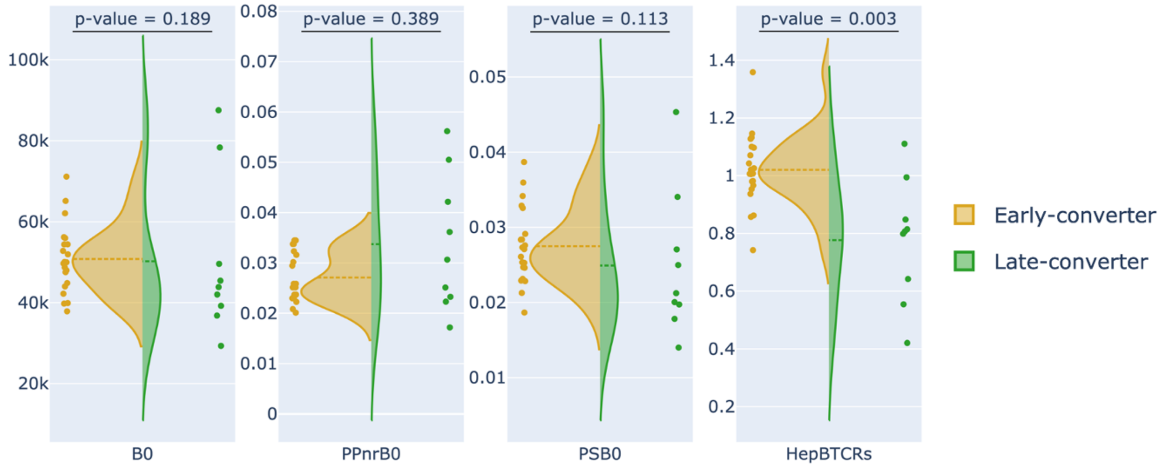


***Additional file 5. CD4+ T cell parameters reveal class differences.*** *Pre-vaccination TCR data feature distribution per class.* HepBTCRs are significantly elevated in early vaccine responses (p-value = 0.003, *two-sided Mann-Whitney U-test*).

*Normalised ratio of vaccine-specific TCRs (*HepBTCRs*); T-Cell Receptor (TCR).*

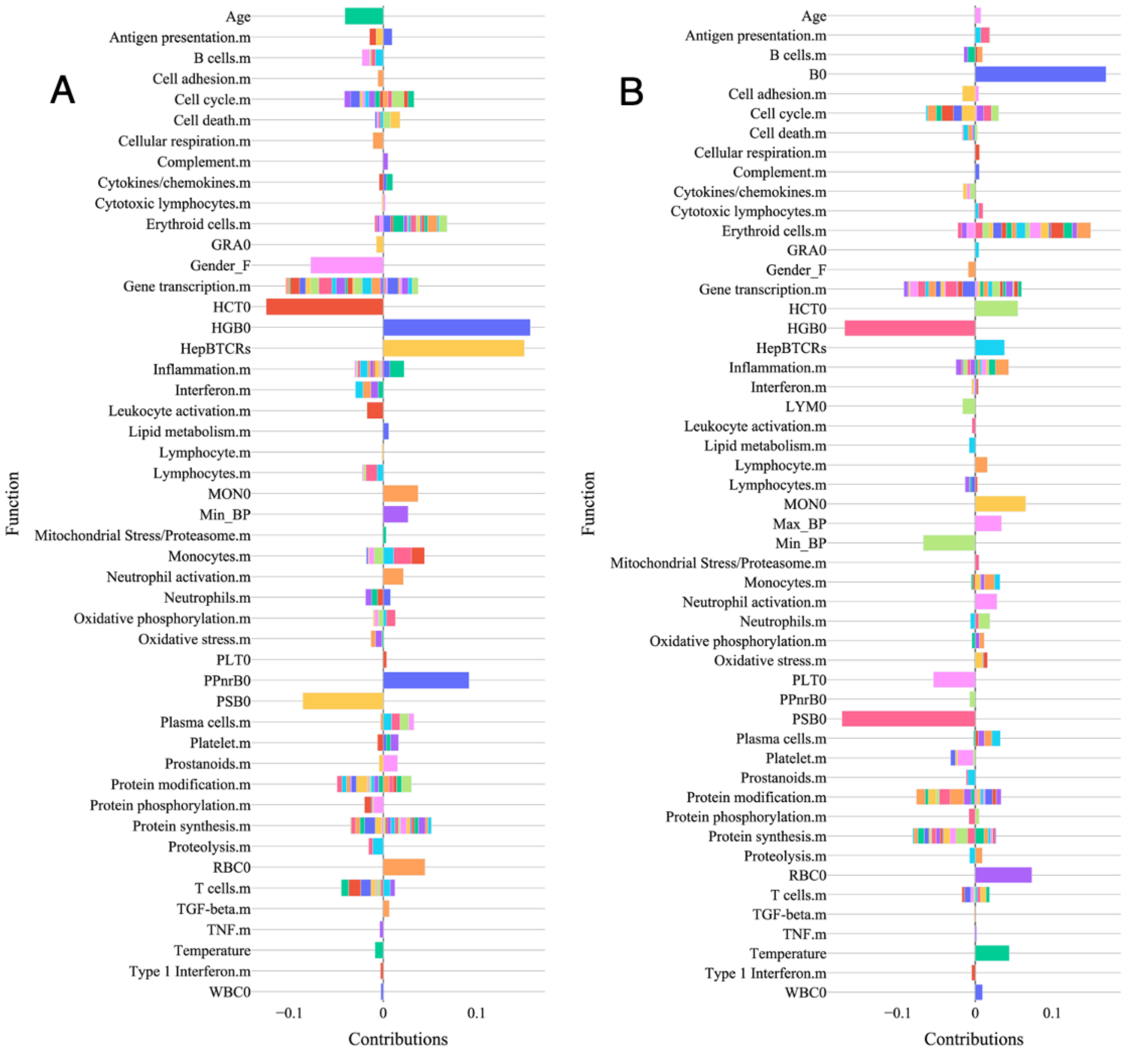


***Additional file 6. MCCA reveals distinct contributions of the metadata and expression.*** *Visualisations of the loadings to the first (A) and second (B) dimensions of the MCCA projection. Coefficients were stacked by function, with null-values and modules without associated functionality being omitted.*

*Multi-view Canonical Correlation Analysis (MCCA).*

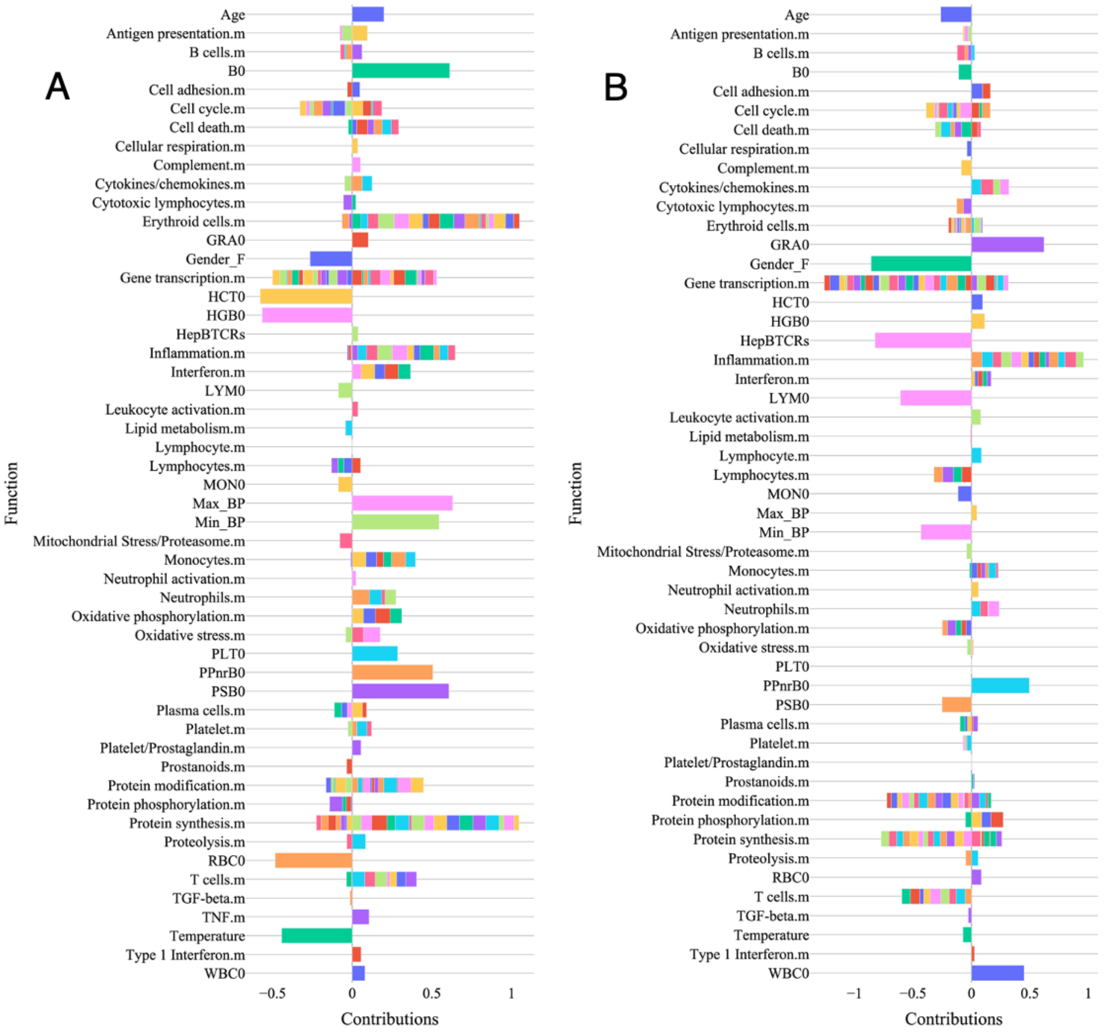


***Additional file 7. GroupPCA components support MCCA findings and offer better interpretability.*** *Visualisations of the loadings to the first (A) and second (B) dimensions of the GroupPCA projection. Coefficients were stacked by function, with null-values and modules without associated functionality being omitted.*

*Multi-view Canonical Correlation Analysis (MCCA); Principal Component Analysis (PCA).*

***Additional file 8. Performance table for the evaluation in terms of AUC and accuracy of a LR trained on single modality data.***

| View | Cell counts | TCR-seq | Metadata | RNA-seq |
|---|---|---|---|---|
| Accuracy | 0.506±0.044 | 0.708±0.067 | 0.598±0.058 | 0.440±0.018 |
| AUC | 0.430±0.038 | 0.731±0.040 | 0.588±0.028 | 0.611±0.009 |

*Area Under Curve (AUC); Logistic Regression (LR).*

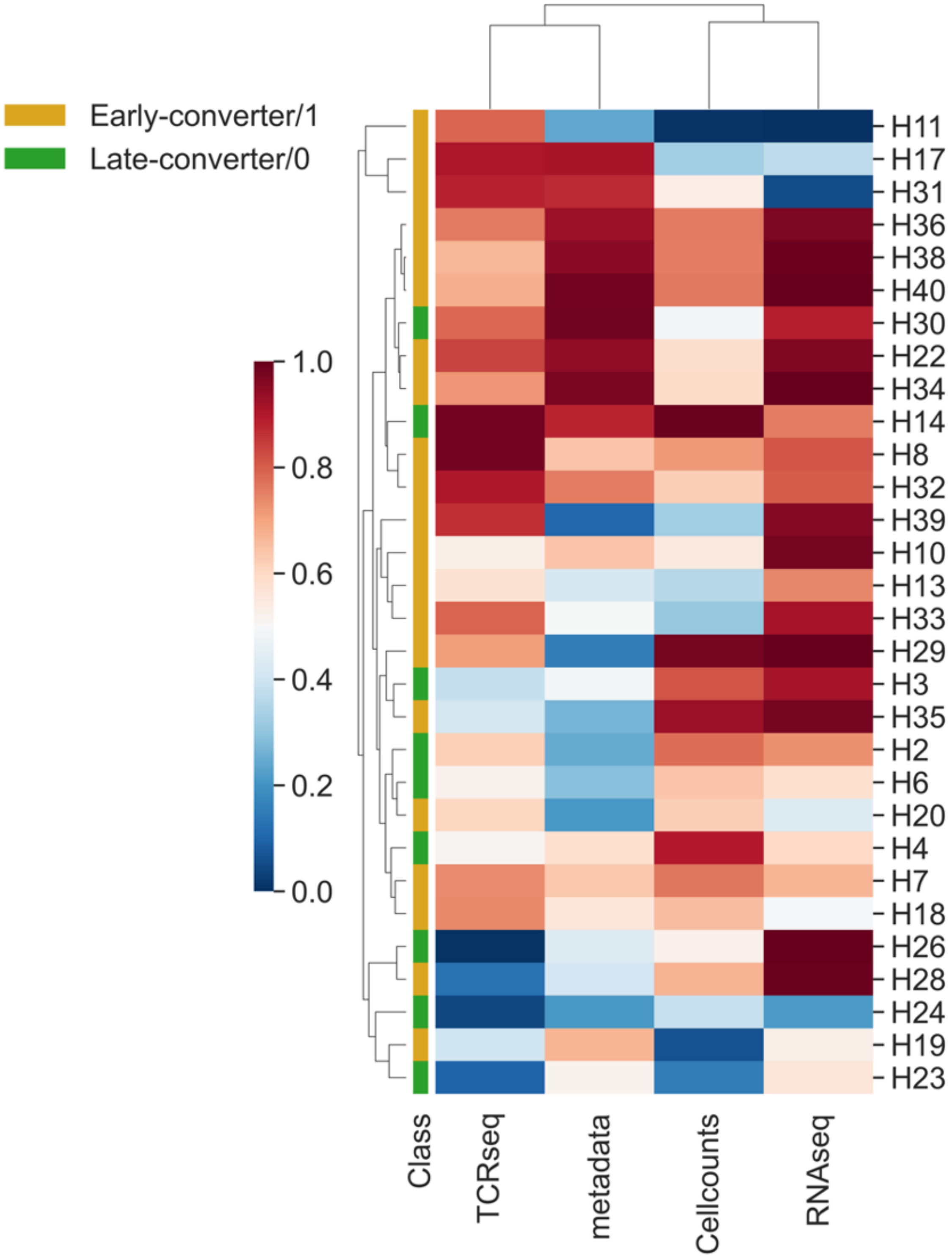


***Additional file 9. Unimodal predicted probabilities show layer to layer difference in predictive power.*** *Predicted probabilities attributed to each patient of the cohort, per single data modality of a LOOCV LR.*

*Leave-One-Out Cross-Validation (LOOCV); Logistic Regression (LR).*
